# Goats Given Transdermal Flunixin Meglumine Displayed Less Pain Behavior After Castration

**DOI:** 10.1101/2020.06.04.134049

**Authors:** Amanda Lee, Meggan Graves, Andrea Lear, Sherry Cox, Marc Caldwell, Peter Krawczel

**Affiliations:** Department of Animal Science, University of Tennessee, Knoxville, USA; Department of Large Animal Clinical Sciences, University of Tennessee, Knoxville, USA; Department ofBiomedical and Diagnostic Sciences, University of Tennessee, Knoxville, USA; Department of Agricultural Sciences University of Helsinki, Helsinki, Finland; Research Centre for Animal Welfare, Department of Production Animal Medicine, University of Helsinki, Helsinki, Finland; Helsinki One Health, University of Helsinki, Helsinki, Finland

## Abstract

Pain management should be utilized with castration to reduce physiological and behavioral changes. Transdermal application of drugs require less animal management and fewer labor risks, which can occur with oral administration or injections. The objective was to determine the effects of transdermal flunixin meglumine on meat goats’ behavior post-castration. Male goats (N = 18; mean body weight ± standard deviation: 26.4 ± 1.6 kg) were housed individually in pens and randomly assigned to 1 of 3 treatments: (1) castrated, dosed with transdermal flunixin meglumine; (2) castrated, dosed with transdermal placebo; and (3) sham castrated, dosed with transdermal flunixin meglumine. Body position, rumination, and head-pressing were observed for 1 h ± 10 minutes twice daily on days −1, 0, 1, 2, and 5 around castration. Each goat was observed once every 5-minutes (scan samples) and reported as percentage of observations. Accelerometers were used to measure standing, lying, and laterality (total time, bouts, and bout duration). A linear mixed model was conducted using GLIMMIX. Fixed effects of treatment, day relative to castration, and treatment*day relative to castration and random effect of date and goat nested within treatment were included. Treatment 1 goats (32.7 ± 2.8%) and treatment 2 goats (32.5 ± 2.8%) ruminated less than treatment 3 goats (47.4 ± 2.8%, *P* = 0.0012). Head pressing was greater on day of castration in treatment 2 goats (*P* < 0.001). Standing bout duration was greatest in treatment 2 goats on day 1 post-castration (*P* < 0.001). Lying bout duration was greatest in treatment 2 goats on day 1 post-castration compared to treatment 1 and treatment 3 goats(*P* < 0.001). Transdermal flunixin meglumine improved goats’ fluidity of movement post-castration and decreased head pressing, indicating a mitigation of pain behavior.

## Introduction

Castration is a common management practice employed in goats to decrease aggression (1–3), reduce tainted meat odor, and prevent off-taste flavor of meat (4). Because castration is often performed without analgesia or anesthesia, the procedure results in pain and distress (2). To date, the use of pain management around castration has not been reported in the USDA NAHMS goat survey (5). Comparatively, on dairy cow operations, 5.6% or less of dairy calves were treated with analgesia or anesthesia during castration (6), suggesting that regardless of species, pain management around castration is minimal in food producing animals. Additionally, only 22% of veterinarian survey respondents indicated they used local anesthetics when castrating beef calves (7). Veterinarians in Canada reported they did not use anesthetics when dairy or beef calves were less than 6 months old or an elastrator was used because of unknown withdrawal periods for opiates, availability of drugs, and the costs of analgesics for cattle, equine, and swine (8). Although castration is well recognized as a painful in numerous species (9–11), anesthetics not often used.

Despite minimal use, veterinarians strongly suggest analgesics are beneficial for the safety of all animals and producers (8). Castration, animal handling, restraint, and incision can increase livestock’s stress, increasing handlers and animals’ risk of injury (12). Among kids disbudded and provided with lidocaine hydrochlorate, injections did not prevent cortisol increases and caused increased frequency and intensity of struggles and vocalizations (13). This indicates that when pain management was provided, it was insufficient to prevent physiological and behavioral changes indicative of pain. Further, castrated calves provided with pain management had lower salivary cortisol concentrations at 4 h post-castration, but no difference at 24 h post-castration, suggesting analgesics stop working before pain has sufficiently dissipated (14). Transdermal dosing provides an alternative means to administer pain mitigation drugs without excessive handling or injections, during stressful procedures (15, 16). This suggests transdermal dosing may be an effective pain-mitigation strategy to use in goats around castration. However, because allogrooming and self-grooming behavior increases in goats after castration (17, 18), transdermal dosing may be challenging to use when goats are group housed and the transdermal drug can be licked or scratched off, versus situations in which goats are individually housed.

Because many animals change their behavior around painful procedures, including castration, one means to assess the efficacy of a transdermal non-steroidal anti-inflammatory drug is through behavioral observations, including lying and standing behavior, feeding, ruminating, grooming, scratching, and head pressing. When provided with isoflurane alone or in combination with other pain management tools, lying and feeding behavior did not vary between groups, suggesting equal pain management between treatments (19). Goats after disbudding struggled more frequently, regardless of pain management strategy, versus goats under stimulated disbudding (13). Cauterized disbudded goats had a greater increase in feeding time post-castration, compared to sham disbudded goats (18). Post-castration, male goats groomed 3 times time more than when intact (17). Goats under toxicology stress were observed head pressing (20), suggesting head pressing can serve as a non-specific measurement of discomfort or pain. This suggests that behavioral changes can be used to evaluate the effects of a transdermal non-steroidal anti-inflammatory in goats using castration as a model for pain treatments.

Although all behaviors were not evaluated in castrated goats, castration in other species can provide further clarity about potential behavioral changes. Calves castrated without pain management displayed more abnormal body postures and decreased willingness to move, compared to calves provided lidocaine and epinephrine anesthesia at 4 and 8 hours post-castration (21). Calves, regardless of age, increased restlessness, tail wagging, foot stamping, and head turning post-castration with rubber rings (22). Among those castrated at 21 or 42 days of age, castrated calves spent less time feeding compared to non-castrated calves (22). This suggests continuous behaviors, including lying, standing, and feeding time and static behaviors, including tail wagging, foot stamping, and grooming, may be effective means to evaluate pain behavior post-castration with transdermal pain management.

Transdermal pain mitigation has the possibility to reduce behavioral changes among meat goats after castration, but has yet to be evaluated in this capacity. Therefore, the objective of this study was to assess the effects of a transdermal formulation of flunixin meglumine (TFM) on meat goats’ behavior after surgical castration, relative to sham-castrated control treated with TFM. Transdermal flunixin meglumine improved goats’ ability to switch body positions post-castration and decreased head pressing, indicating a lessening of pain behavior.

## Materials and methods

All procedures of the current study were approved by the University of Tennessee’s Institutional Animal Care and Use (IACUC #2563-1017). Eighteen intact male goats (mean body weight ± standard error of the mean: 26.4 ± 1.6kg) were enrolled in the study. Prophylactic antibiotics and dewormer were administered to all goats at acquisition. Goats were acclimated to pasture access for 2 weeks and provided with ab-libitum mixed grass hay and goat pellets fed at 0.02% body weight twice daily to facilitate visual observation of acclimation and general health. Two days before study initiation, goats were moved to individually housed pens (range: 2.9 m^2^ to 6.9 m^2^). The minimum requirement for feeder lambs ranging from 14 to 50 kg is 0.74 to 0.93m^2^, indicating that the goats in our study were housed in pens exceeding the minimum requirements for their weight (23). Goats remained within their assigned pen for the duration of the study. After moving to individual pens, goats were fed ad libitum mixed-grass hay from suspended baskets and ad libitum water from buckets. To allow for presentation of natural climbing behaviors (24), goats were provided with a climbing box (0.6 × 0.6 × 0.5m^3^). Mean, maximum, and minimum temperature, relative humidity, and minimum relative humidity were collected at a local weather station (3.2 km away) and mean temperature humidity index (THI) was calculated using the National Oceanic and Atmospheric Administration formula (25). All goats were evaluated by staff from the University of Tennessee, College of Veterinary Medicine, treated with dewormer and metaphylactic antibiotics, and confirmed to be healthy before being enrolled in this study. One goat was removed from the entire data set because of sickness symptoms, unrelated to castration or study, during the entire study, and disease confirmed after study completion.

### Castration treatment

Goats were randomly assigned to one of three treatments, accounting for body weight: castration + transdermal FM (CFM+; mean body weight ± standard error of the mean: 26.02 ± 1.50), castration + placebo transdermal dose (CFM-; mean body weight ± standard error of the mean: 26.10 ± 1.42), and sham castration + TFM (NCFM+; mean body weight ± standard error of the mean: 26.3 ± 1.69). Goats in all groups were sedated using 0.15 mg/kg of xylazine intramuscularly. Animals were laid in right lateral recumbency and the left rear limb of all CFM+ and CFM- goats was elevated and secured with a rope. A subcutaneous ring block of 2% lidocaine was injected around the scrotal neck and the tissue of the spermatic cord. Scrotal skin was aseptically prepared and a standard technique was used to surgically castrate each animal, as described by Ames and Noordsy (26), with a maximum surgery time of 10 minutes. Goats in NCFM+ received a “placebo” castration, in which steps were replicated but not performed to ensure mean procedural time was consistent. A single unblinded individual applied the FM treatment to CFM+ and NCFM+ groups and a “placebo” to the CFM- group. The placebo given was the same color and consistency of TFM, to ensure no visual bias occurred. All other observers and study personnel were blinded to the treatment for the entirety of the study.

### Assessment of behavioral response

#### Visual observation

Goats were observed for 1 hour ± 10 minutes before and after dawn and dusk (04:45 to 06:45 h and 20:30 to 22:30 h) using an ethogram (Table 1), when goats are reported as most active (Zobel, 2019; *Personal Communication*). Observations were made with 5-minute scan samples. Additionally, because the study was conducted during the summer (mean temperature ± standard error of the mean: 23.53 ± 0.65), visual observation around dusk and dawn minimized the influence of heat stress. Visual observation was collected by 3 total observers (1 during each behavioral observation window) on days −1 (before castration), 0 (day of castration), and 1, 2, 3, and 5 (post-castration). One expert scorer with previous training observing goat behavior trained two assistants using an ethogram of goat behavior (Table 1). Each assistant was provided with examples of each behavior through direct observation under the supervision of the lead research technician to confirm all observers understood the behaviors of interest. Interobserver reliability was evaluated by kappa co-efficient (30) in Excel (Microsoft Office, 2019, Albuquerque, New Mexico, USA; κ = 0.79). At dusk two days before castration, the observers visually assessed behaviors using 5-minute instantaneous for 1 hour totaling twelve 5-minute instantaneous samples for 18 goats. For each 5-minute scan sample, observer reliability was evaluated and a total kappa-coefficient was reported.

**Table 1.** Ethogram of goat behaviors recorded during visual observation in which behaviors were recorded every 5 minute from 1 hour before to 1 hour after dawn and dusk (120 minutes total).

| Category | Behavior | Definition |
| --- | --- | --- |
| Body | Climbing | Standing with at least 2 feet on the supplied box [29] |
| Body | Lying | Lying sternally or laterally [34] |
| Body | Standing | Standing idly on all four feet [34] |
| Body | Standing on box | Standing idly on all four feet on the box [34] |
| Body | Lying on box | Lying sternally or laterally on the box [34] |
| Groom | Grooming | Upward scraping motions of the lateral incisor-canine teeth against the pelage and scratch with the hoof of the hindleg, occurs on any area except head and neck [14] |
| Groom | Rubbing | Pushing a body region up or down against a stationary object [34] |
| Mouth | Biting | Taking a new material (non-food and non-self) into the mouth and chewing [35] |
| Mouth | Vocalizing | Vocalizations [36] |
| Mouth | Feeding | Muzzle of goat is inside feeding trough or pile of hay [37] |
| Mouth | Ruminating | Regurgitation of material from the reticulo-rumen to the buccal cavity where solid material is re-masticated and re-insalivated before swallowed [28] |
| Pain | Head Pressing | Animal stands with head-pressed against a surface [38] |

#### Accelerometers

Accelerometers (HOBO pedant G data loggers; Onset Computer Corporation, Bourne, MA, USA) were attached to the lateral side of the right and left hind legs above the metatarsophalangeal using veterinary wrap and elastic adhesive tape. The accelerometers were placed in a vertical position with the x-axis perpendicular to the ground (31). HOBO pendant accelerometers were validated for use in goats to determine lying time and bouts for each side, and standing time and bouts (31).

### Statistical Analysis

Total number of behaviors observed within a day were divided by total number of daily 5-minute scan samples and multiplied by 100% to determine percentage of time that a behavior occurred during the 4-h of daily observation. For all 5-minute scan samples, a body position (lying, standing, climbing on box, standing on box, and lying on box) was recorded. Body position was further summarized into three categories: lying, standing, and on box (climbing, standing and lying on box). An additional oral response (ruminating, feeding, chewing, or vocalizing), grooming behavior (grooming, rubbing, or scratching), or pain behavior (head pressing) was recorded if observed, so that a goat could be recorded with a body and mouth position. For example, a goat could be both lying and grooming or standing and feeding. A linear mixed model was conducted using the GLIMMIX procedure of SAS 9.4 (Cary, NC, USA). Fixed effects of treatment, day relative to castration, and treatment*day relative to castration; covariates of weight and pen size; and random effect of date and goat nested within treatment were included in all models. An auto-regressive structure was used to account for changes over time.

All accelerometer variables (standing, total lying, right lying, and left lying time, bouts, and duration) were summarized by day and assessed for normality before inclusion into final models. A linear mixed model was conducted using the MIXED procedure of SAS 9.4 (Cary, NC, USA). The fixed effects of treatment, day relative to castration, and treatment*day relative to castration; covariates of weight and pen; and random effects of date and goat nested within treatment were included in all models. The lsmeans of non-continuous variables was evaluated to determine treatment differences.

## Results

### Visual observation

Lying percentage and percentage of time on box did not differ by treatment (*P* ≥ 0.63; Table 2). Observed standing percentage was differed by treatment (*P* = 0.04; Table 2) and date (*P* = 0.03; S1 Table). Observed ruminating percentage was different by treatment (*P* = 0.0012) and date (*P* < 0.0001; Table 2). During visual observation, NCFM+ goats spent a greater percentage of time ruminating than CFM+ and CFM- goats (Table 2). Observed feeding time only varied by date (*P* = 0.005; S1 Table), with an increase in observed feeding percentage on days 1 and 2 post-castration, compared to day 0 post-castration for all groups.

**Table 2.** The effects of treatment (castration plus transdermal flunixin meglumine; castration plus transdermal placebo; and sham castration plus transdermal flunixin meglumine) on the percentage of time each behavior was visually observed from 1 hour before to 1 hour after dawn and dusk twice daily on days −1, 0, 1, 2, and 5 relative to castration or sham castration treatment. Different letters indicate differences by treatment (a and b) and differences by day relative to castration (x, y, and z) at *P* ≤ 0.05.

| Behavior | Treatment |  |  |
| --- | --- | --- | --- |
|  | Castration + TFM (%) | Castration + Transdermal Placebo (%) | Sham Castration + TFM (%) |
| Standing | 45.2 ± 5.3<sup>a</sup> | 40.8 ± 5.3<sup>a</sup> | 27.3 ± 5.3<sup>b</sup> |
| Lying | 32.1 ± 8.5 | 39.39 ± 8.5 | 43.9 ± 8.5 |
| On Box | 20.3 ± 6.8 | 22.2 ± 6.8 | 28.9 ± 6.8 |
| Ruminating | 32.7 ± 2.8<sup>b</sup> | 32.5 ± 2.8<sup>b</sup> | 47.4 ± 2.8<sup>a</sup> |
| Feeding | 16.7 ± 2.4 | 17.1 ± 2.4 | 22.4 ± 2.4 |
| Chewing | 0.9 ± 0.3 | 0.3 ± 0.3 | 1.1 ± 0.3 |
| Grooming | 3.8 ± 0.7 | 3.6 ± 0.7 | 4.1 ± 0.7 |
| Scratching | 1.1 ± 0.3 | 1.2 ± 0.3 | 1.4 ± 0.3 |
| Rubbing | 1.5 ± 0.6 | 2.8 ± 0.6 | 1.6 ± 0.6 |

Head pressing was different by treatment and date interaction (Fig 1). Castrated goats not provided flunixin meglumine spent a greater percentage of time head pressing on days 0 and 5 post-castration compared to NCFM+ goats (*P* = 0.03). On the day of castration, CFM- goats spent the most time head pressing, compared to both other treatments (Fig 1). Chewing, grooming, scratching, rubbing, and vocalizing observed percentages did not vary by treatment, and only rubbing was different by date (*P* = 0.02; Table 2).

**Fig 1.**
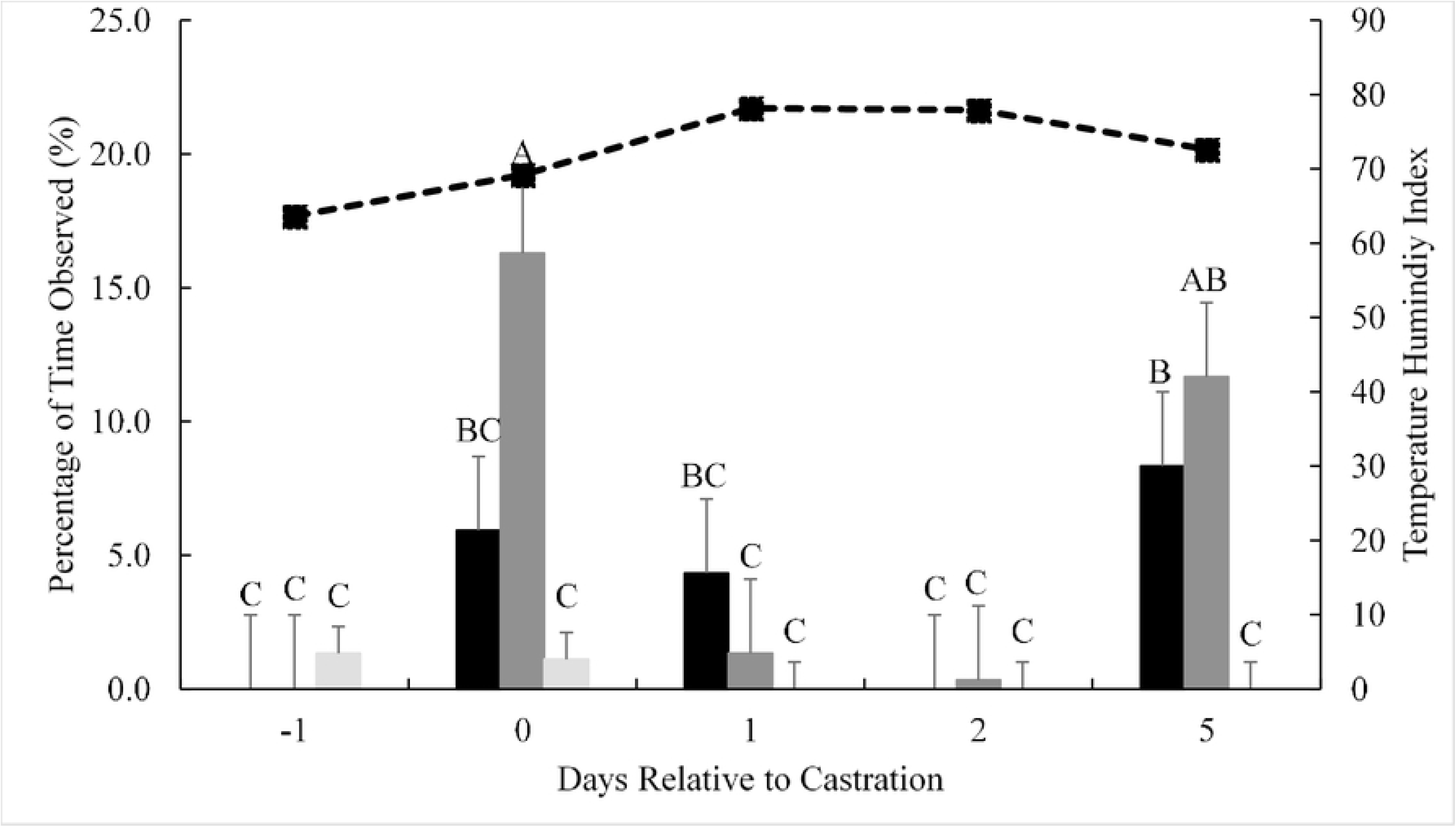
Percentage of time that goats were observed head pressing relative to the treatment and corresponding mean temperature humidity index. ■ = castration plus transdermal flunixin meglumine, 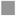 = castration plus transdermal placebo, 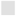 = sham castration with transdermal flunixin meglumine. Different letters indicate differences by treatment x day relative to castration (a, b, c) at *P* ≤ 0.05.

### Accelerometers

Mean standing time, standing bouts, lying time, and lying bouts did not differ with treatment or date (Table 3; *P* ≥ 0.09). Standing bout duration was affected by treatment and date interaction (*P* = 0.03), but was not correlated with THI (Fig 2). On day 1 post-castration, CFM- goats’ standing bout duration was twice as great as any other treatment on any other day (Fig 2). Castrated goats treated with FM and NCFM+ goats tended to spend less total time lying than CFM- goats (*P* = 0.10; Table 2). Left-side lying bout duration and total lying bout duration was different by treatment and date interaction (*P* < 0.001), but was not correlated with THI (Fig 3 and S1 Fig, respectively). Castrated goats with transdermal placebo spent a longer duration lying per bout on the left-side and total on days 1 and 4 post-castration (Fig 3 and S1. Fig, respectively). Regardless of treatment and date, goats spent more time lying on their left-side than their-right side. Mean right-side lying time was 3 to 4 times less per day than mean left-side lying time (Table 3). Left-side lying time tended to be different by treatment, with CFM- goats spending longer lying on their left-side than other treatments (*P* = 0.07; Table 3). All other variables were not affected by treatment or date (*P* ≥ 0.12; Table 3 and S2 Table, respectively).

**Table 3.** The effects of treatment (castration plus transdermal flunixin meglumine; castration plus transdermal placebo; and sham castration plus transdermal flunixin meglumine) on behavior as recorded by an accelerometer (Hobo, Onset, MA, USA; excluding standing bout duration, lying bout duration, and left side lying bout duration) attached to the rear legs of meat goats (N = 17) on days −1, 0, 1, 2, 3, 4, and 5 relative to castration or sham castration treatment.

|  |  | Treatment |  |  |
| --- | --- | --- | --- | --- |
| Behavior | Units | Castration + TFM | Castration + Transdermal Placebo | Sham Castration + TFM |
| Total lying time | Min/24 h | 1068.4 ± 43.9 | 1255.2 ± 69.0 | 1085.8 ± 49.0 |
| Total lying bouts | N | 70.7 ± 13.2 | 37.6 ± 20.6 | 80.7 ± 14.7 |
| Standing time | Min/24 h | 371.6 ± 43.9 | 184.8 ± 69.0 | 354.2 ± 49.0 |
| Standing bouts | N | 16.1 ± 1.6 | 9.6 ± 2.5 | 17.0 ± 1.8 |
| Left lying time | Min/24 h | 817.6 ± 76.2 | 1099.9 ± 119.4 | 693.4 ± 84.9 |
| Left lying bouts | N | 41.9 ± 5.6 | 23.0 ± 9.2 | 42.3 ± 6.5 |
| Right lying time | Min/24 h | 244.7 ± 70.3 | 167.6 ± 110.2 | 393.9 ± 78.4 |
| Right lying bouts | N | 34.0 ± 7.6 | 13.3 ± 13.0 | 37.0 ± 9.2 |
| Right side lying bout duration | Mins/bout | 8.2 ± 2.7 | 16.5 ± 4.2 | 12.0 ± 3.0 |

**Fig. 2.**
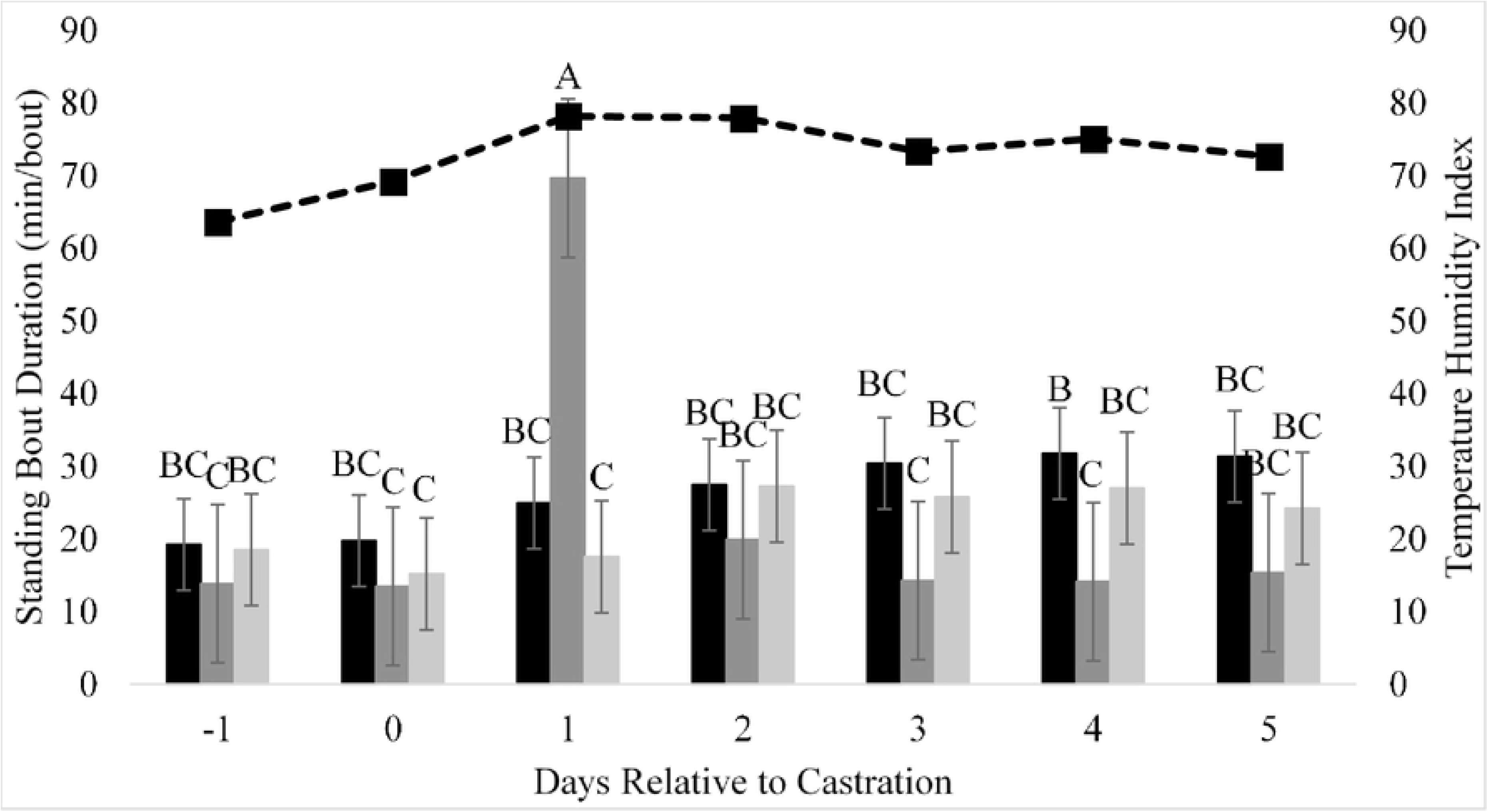
Standing bout duration per day, relative to the day of castration and mean temperature humidity index. ■ = castration plus transdermal flunixin meglumine, 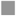 = castration plus transdermal placebo, 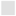 = sham castration with transdermal flunixin meglumine. Different letters indicate differences by treatment x day relative to castration (a, b, c) at *P* ≤ 0.05.

**Fig. 3.**
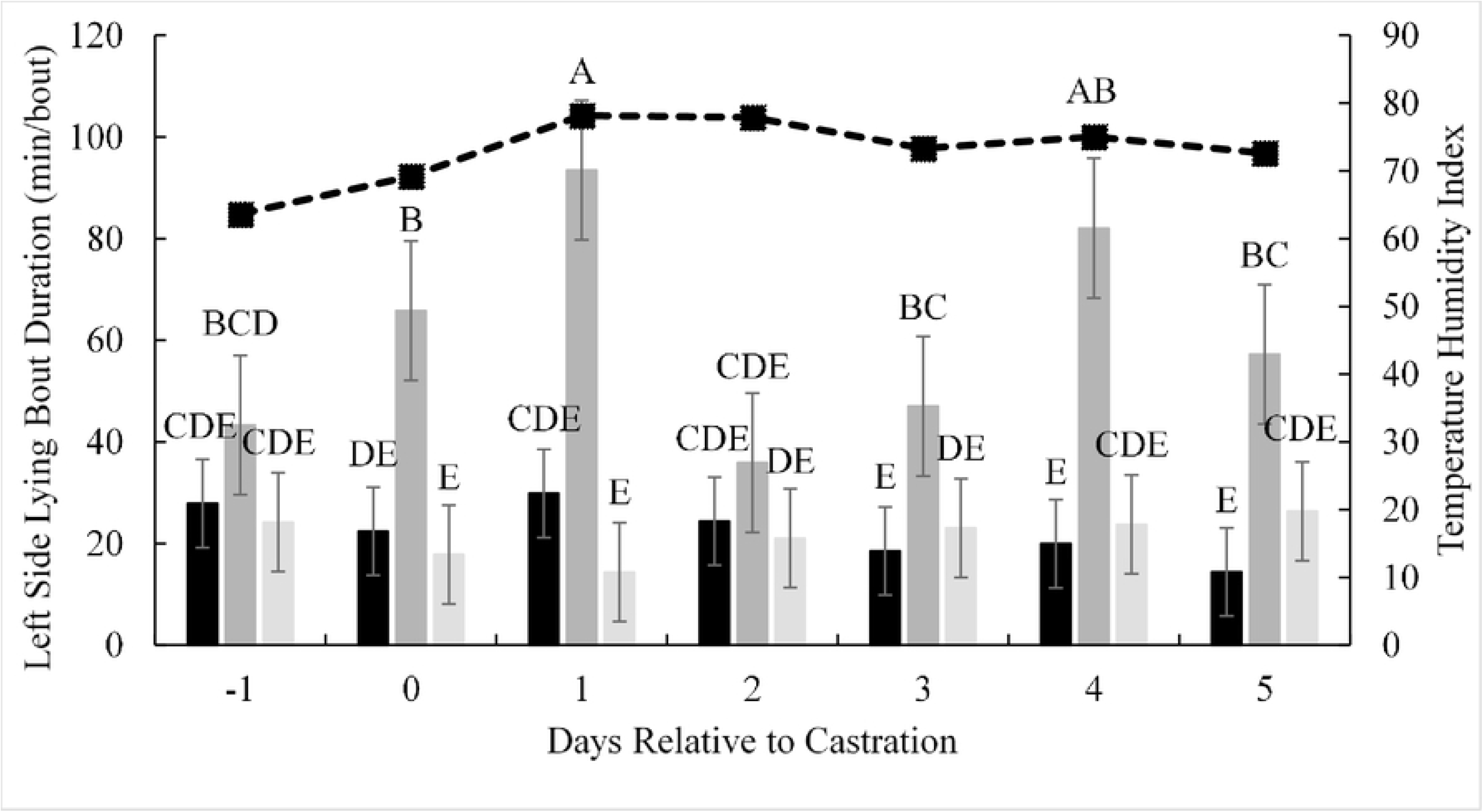
Left side lying bout duration per day (minutes/bout), relative to the day of castration and corresponding mean temperature humidity index. ■ = castration plus transdermal flunixin meglumine, 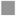 = castration plus transdermal placebo, 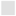 = sham castration with transdermal flunixin meglumine. Different letters indicate differences by treatment x day relative to castration (a, b, c) at *P* ≤ 0.05.

## Discussion

This study evaluated transdermal administration of FM as a means to reduce pain post-castration in meat goats. Providing TFM reduced the amount of time that goats spent head pressing, a non-specific indication of pain. Additionally, goats castrated and given TFM or sham-castrated and given TFM had a shorter standing and lying bout durations, suggesting goats had greater ability to switch between body positions post-castration when receiving TFM.

### Visual observation

Within this study, visual observation was used to assess goat’s body position (lying, standing, or on box), ruminating, feeding, chewing, grooming, scratching, rubbing, and head pressing behavior as an indication of goats’ pain around castration. Although body position did not vary by treatment, goats spent a mean 20 to 29% of their time using the boxes. This suggests that goats recovering from disbudding, will still actively seek out climbing resources. Even within confinement, goats will actively seek opportunities to climb, as climbing is a natural behavior that can help manage hoof growth (24, 32). Additionally, during a 3 h period, goats given saline, lidocaine, and lidocaine and flunixin immediately preceding disbudding had 18.4 to 39.9 exploration incidences, which included climbing (33). This suggests that goats recovering from disbudding, will still actively seek out climbing resources. During our study, all goats were observed using boxes at some time, suggesting box use may not be a sensitive tool to measure comfort following castration, since climbing boxes appeared to be a used resource, regardless of pain.

Rumination time was greater in NCFM+ goats compared to CFM- or CFM+ goats during the 4 h of daily observation. However, on days 1 and 2 post-castration, rumination time overall decreased, suggesting that both treatment groups experienced some pain, driving this change in overall rumination time by date. When lambs were castrated via surgery without pain mitigation, surgery plus flunixin, surgery plus flunixin and lidocaine, or surgery plus lidocaine their mean eating and rumination time (combined as feed intake) did not differ (34). However, in surgically castrated bull calves, rumination score decreased on day one post-castration, so that calves on the day after castration showed less interest in hay or straw compared to other days (35). When individually housed, goats’ mean rumination time ranged from 21.4 to 41.8% of their total day (36), with lambs ranging from 19.4 to 33.3% observed feeding and ruminating during a 12-hour observation period (34). Within our study, observed rumination time dropped by 15 to 20% on days 1 and 2 post-castration, indicating rumination time was cut in half following castration‥ This indicates that overall rumination time and daily rumination time around castration can be important indicators of pain post-castration.

Unlike rumination time, feeding time did not differ between treatments within our study. Although not evaluated in other castration studies, in other pain models such as disbudding, feeding time has also not varied by pain management strategy or species. Among goats disbudded using cautery iron disbudding, and provided with isoflurane gas with and without meloxicam, meloxicam injection alone, oral meloxicam alone, or sham-disbudded, feeding time did not differ (18, 19). Steers surgically castrated at 8 months of age did not differ from bulls or steers surgically castrated at 3 months of age in meal duration, meal eating rate, or daily meals by treatment after surgery (11). In bull calves castrated with Burdizzo, given partial scrotal resection, orchidectomy, or not castrated and provided with xylazine and local sedation, eating time did not differ by treatment (35). This suggests, that feeding time is not sufficiently sensitive to measure pain following castration.

Head pressing, grooming, scratching, and rubbing were assessed as static behaviors to estimate pain or annoyance associated with castration. Although not commonly reported around castration, head pressing has been routinely associated with generalized pain. More than 20% of cases of coenurosis in sheep and pseudorabies in dogs, conditions considered painful, have reported head pressing against a surface (37, 38). Within our study, non-castrated goats spent less than 1% of their time head pressing, whereas castrated goats spent 5 to 7% of their time, suggesting a generalized pain response. Head-pressing increased around castration, and has been anecdotally seen among field veterinarians. Increased head pressing in castrated goats indicates that CFM+ and CFM- goats experienced some pain post-castration, regardless of FM treatment. In future studies assessing pain, head pressing should be explored, as the mechanics and causes of this behavior is not understood outside of head injuries.

### Accelerometers

Within this study, accelerometers were used to assess goats’ lying and standing time, and lying laterality (total time, bouts, and bout duration), as an indication of goats’ comfort around castration. In general, dairy goats housed solely indoors, spent 22% of their time standing (39), which agreed with standing times observed in CFM+ or NCFM+ goats. When considering disbudding as a model for pain behavior, among dairy goats disbudded with cautery iron or sham disbudded, lying time and duration did not differ (18). Dairy goats disbudded via cautery disbudding and provided different pain management methods (isoflurane gas only, meloxicam injection only, meloxicam orally only, or isoflurane gas and meloxicam) also did not differ in lying time or lying duration (19). When using other species to assess castration models, there was similarly no difference in lying or standing behavior. Steers castrated with bands and provided with epidural xylazine plus flunixin meglumine did not differ in lying behavior versus sham castrated steers given epidural xylazine plus flunixin meglumine or sham castrated with saline solution (14). Surgically castrated dairy calves had similar lying and standing times to dairy calves handled but not castrated (22). Beef calves undergoing surgical castration after flunixin meglumine and lidocaine epidural containing epinephrine did not differ in number of steps taken in the 24 h before and post-castration (21). This suggests that lying and standing behavior may not change when animals undergo pain, and is not sufficiently sensitive to assess pain post-castration.

Rather than lying or standing time, lying bout duration may offer a more suitable means to assess comfort related to painful procedures. Within our study, increases in standing bout duration, left-side lying bout duration, and total lying bout duration on the first day post-castration, may suggest that CFM- goats were less comfortable. On average, more than 40% of dairy goats’ lying bout durations were less than 20 minutes/bout, and another 18% (58% total) were less than 40 minutes/bout (31). Within our study, 50% of CFM- goats’ lying bouts were less than 44 minutes/bout, and 62% were less than 52 minutes/bout, while 75% of CFM+ and NCFM+ goats’ lying bouts were less than 30 minutes/bout. This suggests that that although CFM- goats’ mean lying bout duration was longer than the other treatments, lying bout duration was within a normal range. Increases in lying bout durations may indicate that CFM- goats were less comfortable switching positions than CFM+ or NCFM+ goats, as CFM- goats spent longer durations in a single lying bout. Increased lying bout duration has been observed in castration studies conducted in other species. Lambs castrated with conventional rubber rings or novel rubber rings were more restless (had a greater number of switches between standing and lying bouts) than when castrated with novel rubber rings with local anesthetic (10). Bull calves castrated with rubber ring or Burdizzo plus rubber ring shifted between positions more in the three h post-castration, compared to sham handled, Burdizzo alone, or surgically castrated bull calves (9). In our study, increased standing and lying bout duration on the day after castration, indicated CFM- goats had difficulty becoming comfortable, and, therefore, did not want to change between standing and lying as frequently. As shifting positions is seen under multiple castration methods and species, lying bout duration may serve as a suitable and responsive measurement for discomfort around castration.

## Conclusions

Regardless of pain management strategy, post-castration, animals change their behavior to compensate for experienced pain. Providing TFM in addition to lidocaine reduced pain behavior, such as increased lying and standing bout duration on day 1 post-castration, maintaining the percentage of time spent ruminating in NCFM+ goats, and reducing incidence of head-pressing in CFM+ goats post-castration. Understanding how meat goats’ behavior changes around castration can help to determine how effective TFM is at minimizing pain. Within this study, rumination time, head pressing, and lying position served as important indicators of pain post-castration in meat goats. Transdermal flunixin meglumine improved goats’ ability to move between postures on the day following castration and decreased head pressing, indicating TFM use can help to mitigate pain behavior.

## Acknowledgements

The researchers thank the funding and support of the USDA NIFA Animal Health Project (Award # NI18AHDRXXXXG049). Thank you to Dr. Liesel Schnieder for statistical consulting. Thank you to Austin Turner and VTREC staff for helping to feed and take care of the goats during this trial. Thank you to Megan Wright, Alex Shanks, Nicole Harris, Ashley Campeaux, and Desmond Coates for their help with sample collection. Thank you to Danielle Fannon and Jessica Jaggers for their help with behavioral assessment and technology application. Thank you to Ray Ellison for the construction of the climbing boxes for each goat.

## Supporting Information

**S1 Fig. Total bout duration per day (minutes/bout), relative to the day of castration and corresponding mean temperature humidity index.** ■ **= castration plus transdermal flunixin meglumine,** 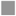 **= castration plus transdermal placebo,** 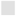 **= sham castration with transdermal flunixin meglumine.**

**S1 Table. The effects of day relative to castration (days −1, 0, 1, 2, 3, 4, and 5) on the percentage of time each behavior was visually observed from 1 hour before to 1 hour after dawn and dusk twice daily on days −1, 0, 1, 2, and 5.** Different letters indicate differences by treatment x day relative to castration (a, b, c) at *P* ≤ 0.05.

**S2 Table. The effects of day relative to castration (days −1, 0, 1, 2, 3, 4, and 5) on behavior as recorded by an accelerometer (Hobo, Onset, MA, USA; excluding standing bout duration, lying bout duration, and left side lying bout duration) attached to the rear legs of meat goats (N = 17).**

